# Condensin II inactivation in interphase does not affect chromatin folding or gene expression

**DOI:** 10.1101/437459

**Authors:** Nezar Abdennur, Wibke Schwarzer, Aleksandra Pekowska, Indra Alon Shaltiel, Wolfgang Huber, Christian H Haering, Leonid Mirny, François Spitz

## Abstract

Condensin complexes have been proposed to play a prominent role in interphase chromatin organization and control of gene expression. Here, we report that the deletion of the central condensin II kleisin subunit *Ncaph2* in differentiated mouse hepatocytes does not lead to significant changes in chromosome organization or in gene expression. Both observations challenge current views that implicate condensin in interphase chromosomal domain formation and in enhancer-promoter interactions. Instead, we suggest that the previously reported effects of condensin perturbation may result from their structural role during mitosis, which might indirectly impact the re-establishment of interphase chromosomal architecture after cell division.

## Introduction

Metazoan chromosomes fold into highly diverse and dynamic conformations constrained by organizational processes that act across genomic length scales (Dekker and Mirny, 2016; Dixon et al., 2016; Gibcus and Dekker, 2013). Chromosomal architectures differ drastically between interphase and mitosis. In interphase, Hi-C maps of interphase genomes have revealed distinct cell-type and locus-specific patterns of chromatin contact frequency. These include sequences of alternating intervals that indicate compartmental segregation of active and inactive chromatin (Lieberman-Aiden et al., 2009), topologically associating domains (TADs), which reflect local compaction (Dixon et al., 2012; Nora et al., 2012), and TAD-associated peaks (Rao et al., 2014) and tracks (Fudenberg et al., 2016) of enriched contact frequency between pairs of loci in *cis*. As cells progress through mitosis, these prominent interphase features disappear as chromosomes get compacted into dense arrays of chromatin loops (Nagano et al., 2017; Naumova et al., 2013). These differences in chromosome organization are tightly related to changes in genomic activities, notably gene regulation in interphase and chromosome segregation during mitosis.

The structural maintenance of chromosomes (SMC) complexes, cohesin and condensin, have been implicated as essential players in the organization of distinct aspects of chromosome organization (Gause et al., 2008; Hirano, 2016; Merkenschlager and Nora, 2016; Rana and Bosco, 2017; van Ruiten and Rowland, 2018). While these two related complexes were originally identified as key determinants of proper segregation of chromosomes during mitosis, a mounting body of evidence has accumulated for their role in the organization of interphase chromosomes (reviewed in (Hirano, 2016; Rana and Bosco, 2017; Yuen and Gerton, 2018)). We and others recently demonstrated that cohesin complexes are essential for the appearance of TADs and their associated patterns (e.g., tracks and peaks) in Hi-C maps (Haarhuis et al., 2017; Rao et al., 2017; Schwarzer et al., 2017; Wutz et al., 2017). The consequences of altered of cohesin dynamics (through inactivation of its loading/unloading factors *Nipbl*, *Mau2* and *Wapl*) (Haarhuis et al., 2017; Schwarzer et al., 2017) are consistent with the TADs resulting from a loop extrusion mechanism (Fudenberg et al., 2016; Sanborn et al., 2015) (reviewed in (Fudenberg et al., 2018)). Importantly, this cohesin-dependent process is not required for the segregation of active and inactive chromatin into compartments (Rao et al., 2017; Schwarzer et al., 2017; Wutz et al., 2017). Rather, cohesin-mediated loop extrusion interferes with chromatin compartmentalization (Nuebler et al., 2018) showing that TADs and compartments are organized by different principles and mechanisms (Rao et al., 2017; Schwarzer et al., 2017).

While cohesin’s involvement in the organization of the interphase genome architecture has by now been clearly established, the function of condensin complexes in the formation of TADs and compartments remains unclear. Most metazoans have two types of condensin complexes, which are composed of the same SMC2-SMC4 dimers that associate with distinct non-SMC subunits (Ono et al., 2003). Condensin I and II serve different functions in the compaction of mitotic chromosomes (Green et al., 2012; Ono et al., 2003). During interphase, only condensin II is reportedly present in nuclei, whereas condensin I is sequestered in the cytoplasm (Gerlich et al., 2006; Hirota, 2004; Ono et al., 2004), although recent data indicate that a small fraction of condensin I may be also be in the nucleus during interphase (Dowen et al., 2013; Li et al., 2015; Zhang et al., 2016). In *C. elegans*, a distinct condensin complex (condensin I^DC^) has been described and shown to be essential for the organization of TAD-like domains in hermaphrodite X chromosomes (Crane et al., 2015). This overall structure leads to a chromosome-wide reduction in gene expression that contributes to balancing expression levels between XX hermaphrodite and X0 male animals (Crane et al., 2015; Jans et al., 2009).

In other species, experimental evidence suggested that condensin II contributes to interphase chromosomal organization and gene expression. In *Drosophila*, mutations in the condensin II kleisin subunit *Ncaph2* reduce the axial compaction of chromosomes, which is thought to be required for favoring intra- over inter-chromosomal interactions and the establishment of chromosome territories (Bauer et al., 2012). ChIP-seq studies reported co-localization of condensin II with other architectural and insulator proteins at the borders of *Drosophila* domains, suggesting that it could contribute to domain boundary formation or strength (Van Bortle et al., 2014). This pattern, which is mediated by interaction with TFIIIC, appears to be conserved in mice and humans (Yuen et al., 2017). Supporting the notion that the binding of condensin II to chromosomes contributes to gene regulation, knockdown of *NCAPH2* was proposed to lead to down-regulation of genes close to TAD boundaries (Yuen et al., 2017). Several other reports provided circumstantial evidence implicating condensin I and II in the control of gene expression (Kranz et al., 2013; Schuster et al., 2013; Woodward et al., 2016), although the underlying mechanisms might have been indirect in most of these cases. Noteworthy, recent studies suggested the recruitment of condensin complexes to gene-regulatory elements, notably super-enhancers (Dowen et al., 2013; Li et al., 2015), where they were proposed to contribute to enhancer activation and looping interaction with promoters (Li et al., 2015) and sustained expression of cell-identity genes (Dowen et al., 2013). However, the experimental setup used in the later studies (either dividing embryonic stem (ES) cells or interference performed as cells are stripped from serum but not yet fully blocked in G0/G1) does not exclude the possibility that the observed changes are indirect consequences of mitotic defects that resulted in aberrant chromosome organization upon exit from mitosis.

To address the uncertainty regarding the role of the condensin II complex in interphase genome organization and gene expression, we deleted the central condensin II kleisin subunit *Ncaph2* in post-mitotic adult mouse hepatocytes. Remarkably, this deletion did not result in any measureable alterations in chromatin organization, as compared to wild-type and *Nipbl* deletions as negative and positive controls, respectively. TADs and compartments appear unaffected. The expression level of only a handful of genes show significant changes and most of those genes are functionally related to mitosis. Our data suggest that, in contrast to cohesin, condensin II is dispensable for the organization of chromosomal domains in non-dividing cells and that its direct contribution in the control of interphase gene expression is negligible.

## Results and Discussion

To determine the functions of condensin II complexes in interphase chromatin folding and gene expression, we deleted critical exons of the gene encoding the *Ncaph2* kleisin subunits (**figure 1A**), which is essential for the activity of the condensin II complex during mitosis. Homozygous deletion of *Ncaph2* is lethal in mice and leads to rapid death of proliferating cells (Nishide and Hirano, 2014). To avoid confounding effects, we used a tissue-specific conditional strategy to induce acute deletion of the gene in non-dividing cells. We achieved this by combining a conditional floxed allele *Ncaph2*^*tm1a(EUCOMM)Wtsi*^(Skarnes et al., 2011), hereafter referred to as *Ncaph2*^*flox*^, with a liver-specific tamoxifen-inducible CRE driver *Tg(Ttr-cre/Esr1*)*^*1Vco*^(Tannour-Louet et al., 2002). We previously used the same strategy to successfully delete the cohesin loading factor *Nipbl* (Schwarzer et al., 2017) and thereby demonstrated its requirement for TAD formation. We followed the same procedure for *Ncaph2*. Ten days after tamoxifen injections, we collected hepatocytes from *Ncaph2*^*flox/flox*^; *Tg(Ttr-cre/Esr1*)*^*1Vco*^ and control animals (**figure 1B**). *Ncaph2* deletion was extremely efficient as judged by mRNA and protein level quantification (**figure 1C-F**).

**Figure 1:**
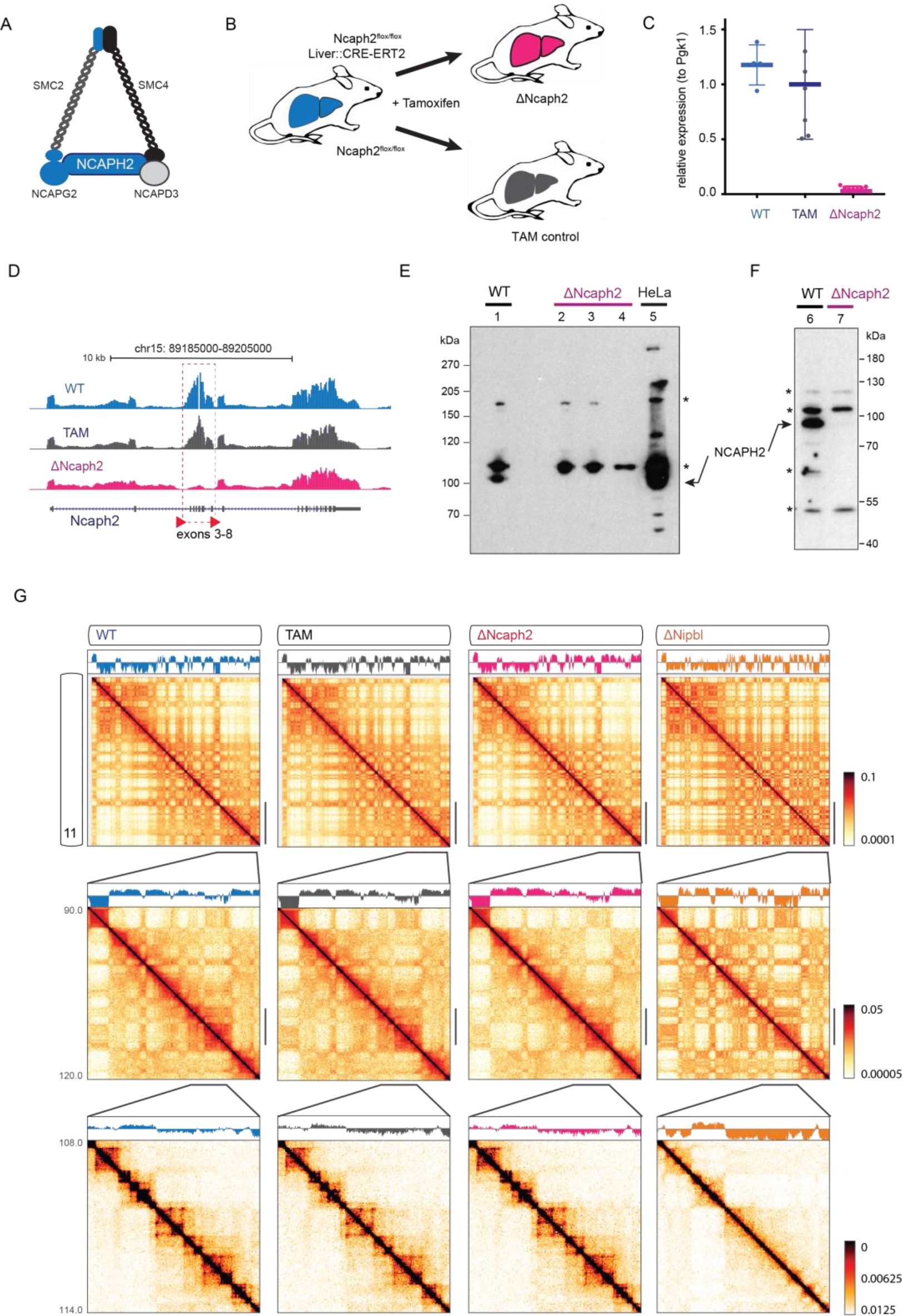
Experimental Strategy. **(A)**Schematic representation of the condensin II complex and its subunits. **(B)**Different groups of mice analyzed. Animals with ΔNcaph2 hepatocytes were produced by intraperitoneal injection of tamoxifen in *Ncaph2*^*flox/flox*^, *Tg(Ttr-cre/Esr1*)*^*1Vco*^ adult mice. Two different controls were used, either untreated (U) animals (genotype: *Ncaph2*^*flox/flox*^, *Tg(Ttr-cre/Esr1*)*^*1Vco*^) or TAM animals that received the same tamoxifen treatment but were not carrying the *Cre* transgene and therefore retained a functional *Ncaph2*^*flox/flox*^ genotype. **(C)**Expression level of *Ncaph2* in hepatocytes. Each dot shows an independent measurement of normalized RT-qPCR expression. Whiskers indicate standard deviation. Expression levels of *Ncaph2* in TAM and ΔNcaph2 are significantly different (TAM: 1.000 ± 0.1887 s.e.m.; ΔNcaph2: 0.05603 ± 0.005896, n=7; p-value = 0.0003 (unpaired two tailed *t*-test). **(D)**RNA-Seq tracks on the *Ncaph2* locus show deletion of the three floxed exons in ΔNcaph2 hepatocytes. Data pooled from 4 independently processed animals for each group. **(E-F)**Western blot detection of NCAPH2 in ΔNcaph2 and control samples. **(G)**Hi-C maps of chromosome 11 across four conditions with compartment eigenvector tracks plotted above. From left to right: WT, TAM, ΔNcaph2 and ΔNipbl. The top row shows whole chromosome maps, the middle row shows a zoom-in of a 30-Mb region, and the bottom row shows a further zoom-in of a 6-Mb region. The color mapping in the top and middle row is logarithmic to highlight differences in compartmentalization, and linear in the bottom row to highlight TAD-associated features.

To examine the effects of condensin II knockout on genome organization, we performed tethered chromatin conformation capture (Kalhor et al., 2012), referred below as Hi-C. For each condition, we produced two different biological replicates (**table S1**). The data from replicates showed high degree of correlation (**figure 1E,F, S2A, S3A**) and we therefore pooled them in subsequent analyses. We compared key features of Hi-C maps, including the global scaling of contact frequency *P(s)*, compartmentalization, contact insulation, contact frequency peaks, and aggregate maps of genomic landmarks associated with chromatin architecture. Unlike the dramatic effects that result from the loss of chromatin-associated cohesin in ΔNipbl (Schwarzer et al., 2017), we observed no substantial changes upon inactivation of condensin II. Examination of Hi-C maps did not reveal alterations of TAD or compartmentalization patterns (**figure 1G**).

To analyze the contact maps of condensin II-inactivated cells in more detail, we calculated the curves of the contact probability *P(s)* as a function of genomic separation *s*. The proximal shallow region of the *P*(*s*) curve (*s* ≈ 100−300 kb), referred to as the shoulder (Fudenberg et al., 2018), reflects local compaction. This region is sensitive to the loss of cohesin, as seen across several studies, consistent with the loss of compaction by loop extruding cohesins. By contrast, the shoulder of *P(s)* cells did not differ between hepatocytes from ΔNcaph2, WT or tamoxifen-injected (TAM) control animals (**figure 2A, S1A**). We also failed to identify any apparent difference for *P(s)* restricted to locus pairs within and between wildtype domain intervals (mostly TADs) and within and between A/B compartmental intervals (**figure S1B,C**).

**Figure 2:**
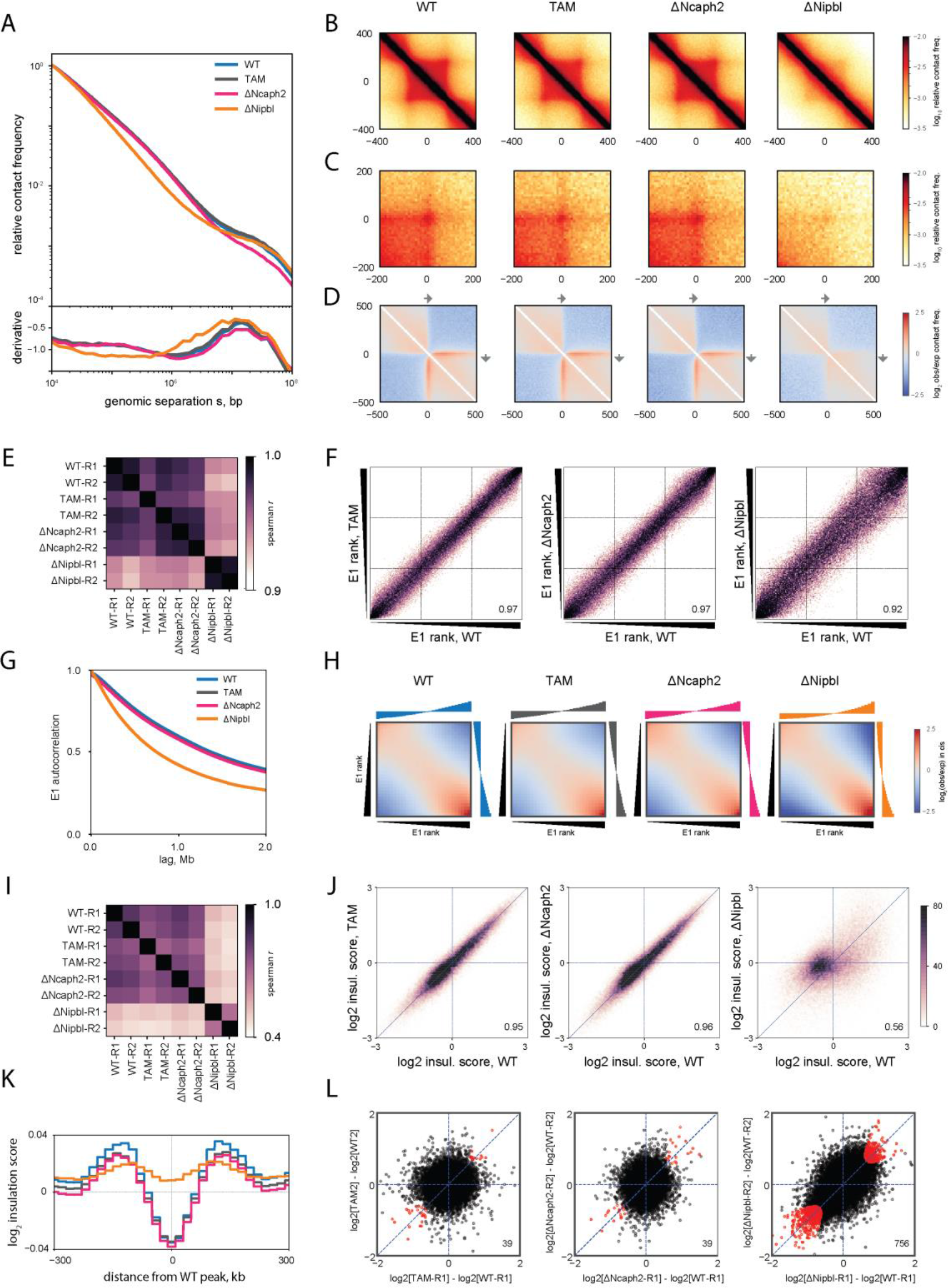
Condensin II removal in differentiated hepatocytes does not significantly impact interphase chromatin organization. **(A)**Top panel: genome-wide P(s) contact frequency curves in WT, TAM control, ΔNcaph2 and ΔNipbl. Bottom panel: derivatives of P(s) curves. **(B, C, D)**Average Hi-C maps centered at **(B)**WT domains of length 300-400kb, **(C**) 102 Hi-C peaks with separation 500-600kb, **(D)**20,000 CTCF binding sites detected in WT from (Schwarzer et al., 2017), oriented by the core CTCF motif. **(E)**Heatmap of Spearman correlation coefficients of compartment tracks among replicates. **(F)**Rank-based scatter plots of compartment eigenvectors between pairs of conditions: TAM, ΔNcaph2, and ΔNipbl versus WT at 20-kb resolution. *r* values inset. **(G)**Genome-wide autocorrelation function of the compartment eigenvectors. **(H)**“Saddle” heatmap plots of compartment signal for all four conditions. Each element of the heatmap contains the average observed-over-expected contact frequency between pairs of loci (20-kb bins) arranged by rank of their compartment score (E1 rank) and grouped into 30 quantiles. The histograms along the top and right axes show the median compartment scores associated with each quantile of genomic bins. **(I)**Heatmap of Spearman correlation coefficients of log2 insulation score tracks among replicates. **(J)**Scatter plots of log2 diamond insulation score between pairs of conditions: TAM, ΔNcaph2, and ΔNipbl versus WT at 20-kb resolution. Spearman *r* values inset. **(K)**Profiles of average insulation score at called insulating loci called in WT. **(L)**Scatter plots of log_2_ fold change in insulation score for each of TAM, ΔNcaph2, and ΔNipbl relative to WT at loci determined to be insulating in at least one condition. Points corresponding to significant differentially insulated loci are colored red.

The most striking effect of cohesin removal is the ubiquitous loss of the features that collectively make up mammalian TADs (local enrichment/insulation, tracks, corner peaks, (Fudenberg et al., 2018)). In contrast with ΔNipbl (Schwarzer et al., 2017), average maps of TADs in ΔNcaph2 do not differ from those of controls (**figure 2B**). Similarly, no effect is observed in average Hi-C peak maps, where ΔNipbl resulted in a fourfold reduction in contact frequency, nor in the asymmetric average contact footprint of CTCF sites which disappears in ΔNipbl (**figure 2C,D**).

Next, we assessed changes in A/B compartmentalization, which reflects segregation of active and inactive chromatin, and is manifested as genome-wide checkerboard patterns on Hi-C maps (Imakaev et al., 2012; Lieberman-Aiden et al., 2009) (**figure 2E,F**). The strength of compartmentalization was increased in studies of acute cohesin degradation (Rao et al., 2017; Wutz et al., 2017), and in the case of persistent cohesin removal via *Nipbl* deletion in postmitotic liver cells, we additionally observed the emergence of a finer compartmental segmentation in large gene-dense regions, quantifiable as a steeper decay of the compartment signal’s autocorrelation function (Schwarzer et al., 2017). In sharp contrast, *Ncaph2* deletion did not result in any of these effects (**figure 2G,H**). Indeed, the ΔNcaph2 and control eigenvectors were strongly correlated (**figure 2E,F**) and there was no change in the autocorrelation function of the eigenvector compared with controls (**figure 2G**). Taken together these results show that condensin appears to play no measurable role in the formation of TADs and associated features, nor in compartmental segregation of active and inactive chromatin in interphase.

Since the loss of chromatin-associated cohesin resulted in genome-wide changes in the diamond insulation score profile, we also examined changes in contact insulation (**figure 2I,J**) in *Ncaph2* mutants. The minima of the insulation score profile correspond to loci for which the contact frequency between neighboring loci on opposite sides is relatively depleted - typically the boundaries of TADs and of compartmental intervals (Crane et al., 2015). The loss of TADs in ΔNipbl cells led to dramatic changes in the insulation profile, as it can be observed in the scatter plot of genome-wide insulation scores against WT. However, the ΔNcaph2 insulation profile correlates as strongly with WT as does that of the TAM control condition. The lack of effect of *Ncaph2* deletion was further highlighted in the depth of the insulation score “valley” averaged around insulating loci (boundaries) identified in WT, which was preserved in TAM and ΔNCaph2 but raised in ΔNipbl (**figure 2K**).

We further tested whether *Ncaph2* deletion may have caused more fine-grained specific changes in local insulation. To detect differentially insulating (DI) loci we used an approach similar to that employed to detect differentially expressed genes (see Methods). For comparison, we applied the same analysis on ΔNipbl cells (having an established effect) and the TAM control cells versus WT. Using the same stringency cutoff, we detected 752 insulating loci as having a significantly different insulation score in ΔNipbl cells vs WT, while only 39 of loci cells were differentially insulating in ΔNcaph2 against WT (**figure 2L**). We found also 39 differentially insulating loci when we compared TAM against WT, and a similar number was obtained when we apply the same procedure on control comparisons of insulation changes between replicates within the same condition (**figure S3**). This implies that the detectable DI loci upon condensin loss are largely false positives and not significant. We conclude that local chromosome organization remains unchanged upon loss of condensin II in interphase.

To test for more subtle or specific effects, we focused on specific regions that have been previously associated with condensin II function. Condensin II has been implicated in certain genomic regions including (1) histone gene clusters, (2) super-enhancers, in mouse ES cells, (3) TFIIIC and tRNA genes, in *Schizosaccharomyces pombe* and mammals. For each of these features we examine changes in interaction frequencies between the loci. We first examine the interaction between the two histone clusters present on chromosome 13, which was reported to depend on condensin II (Yuen et al., 2017). We indeed detect particularly strong interaction between the two clusters, but did not note differences between ΔNcaph2 and controls (**figure 3A**). We then examined interactions between super enhancers (Pott and Lieb, 2014), for which we considered a curated list for mouse liver (Khan and Zhang, 2016) (**figure S4**) or identified such regions *de novo* by locally stitching mouse liver H3K27ac sites as in (Hnisz et al., 2013) (**figure 3B**). We saw little enrichment of enhancer-enhancer interactions above the level observed for regions of the same expression level and hence explained by compartmentalization of euchromatin. In all cases, there was no significant change in enrichment between ΔNcaph2 and controls, while a mild increase in pairwise contact frequency could be detected in ΔNipbl, consistent with increased compartmental segregation upon cohesin loss (Rao et al., 2017; Schwarzer et al., 2017). We then examined the role of condensin in organizing RNAP-III-associated regions. We produce average contact profiles centered at paired POLR3D (RPC4) (**figure 3C**) and POLR3A (RPC1) (**figure S5**) binding sites identified by ChIP-seq in mouse liver (Canella et al., 2012) as well as at the locations of TFIIIC ChIP-seq sites identified in mouse ES cells (Yuen et al., 2017) (**figure S5**). Again, we saw no significant change in enrichment between ΔNcaph2 and controls. Finally, we examined the average Hi-C map centered at the coordinates of pairs of convergently oriented CTCF sites: contact enrichment was lost in ΔNipbl but preserved in ΔNcaph2 (**figure 3D**), just as it was for Hi-C peak coordinates.

**Figure 3:**
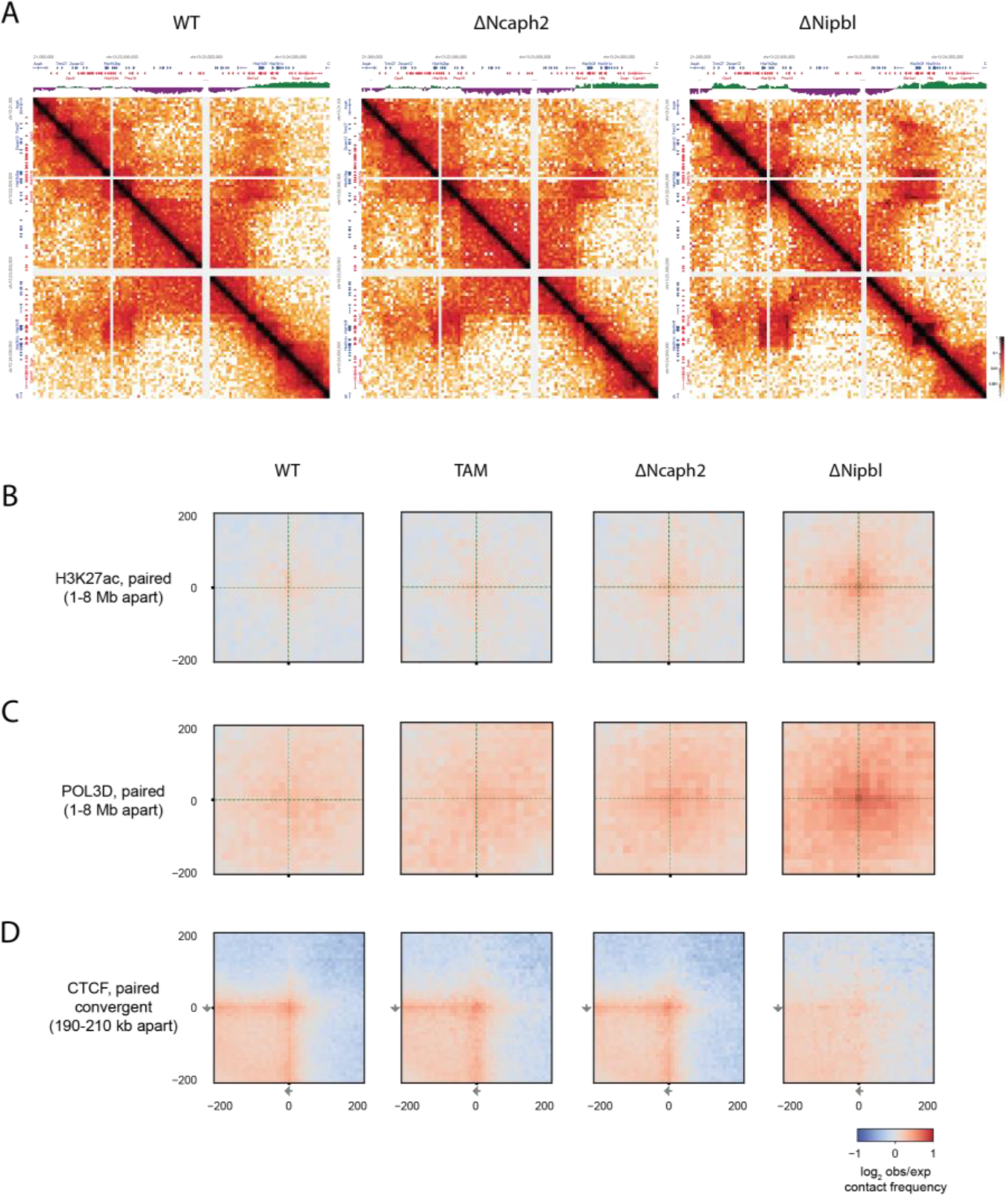
Condensin II removal does not influence specific interactions in the Hist3 gene locus, nor those between transcriptionally active or insulating genomic landmarks. (A) Hi-C maps of the histone gene cluster locus on mouse chromosome 13 in WT (left), ΔNcaph2 (middle) and ΔNipbl (right) displayed in HiGlass (Kerpedjiev et al., 2018). Gene annotations and compartment eigenvector tracks are displayed along the borders. (B, C) Average observed/expected Hi-C pileup maps around the interaction points of paired H3K27ac peaks and Pol3 peaks with separations in the range of 1-8 Mb in *cis*. (D) Average observed/expected Hi-C pileup maps around the interaction points of paired CTCF sites with separations of 190-210 kb in *cis*.

In brief, at the level of resolution and depth offered by our Hi-C data (10-20 kb, 52-75 million valid pairs in *cis* at 10-kb separation or greater, on par with that of other Hi-C of primary tissues (Schmitt et al., 2016)), we did not identify significant differences between the structural partitioning of normal and condensin II-deficient chromosomes. Even though we cannot formally exclude a role in features that are difficult to assess with Hi-C (e.g., spatial positioning within the nucleus (Falk et al., 2018) or nucleus size, (Fazzio and Panning, 2010; George et al., 2014)) or in highly repetitive regions (e.g., centromeres, pericentric regions), we conclude that condensin is involved neither generally nor specifically in the maintenance of interactions that impinge upon most chromosomal loci.

We next investigated if the absence of *Ncaph2* led to gene expression changes (**figure 4**). For this, we compared the transcriptome of control TAM and ΔNcaph2 liver hepatocytes (4 biological replicates per condition). A statistical analysis with DESeq2 (Love et al., 2014) showed that only a very small number of genes were differentially expressed between TAM and *Ncaph2*-deficient hepatocytes (**table 1)**. Gene ontology analysis of these 64 genes showed strong enrichment for genes associated with cell-cycle and cytokinesis (**table 2**). About 20 genes that had been reported to show expression changes during cell cycle (predominantly up-regulated in G2/M) were up-regulated in ΔNcaph2 cells. This coherent shift most likely represents a cell-cycle defect in the few hepatocytes that underwent through mitosis during the course of the experiment in order to maintain liver homeostasis, as knockdown of condensin II has been shown to lead to extended prometaphase (Green et al., 2012). Most of the other mis-regulated genes have been shown to exhibit a circadian pattern of expression (Fang et al., 2014; Wang et al., 2018). The predominance of circadian genes (defined as gene showing an expression profile varying with at least a 2-fold amplitude during a 24-h period) is highly significant (two-tailed Chi-square test with Yates’ correction, *P*<0.0001). However, the correlation between down/up-regulation and normal peak period of the genes is consistent with a shift of a few hours between the two groups of animals and therefore is more likely to correspond to differences in time of sample collection, rather than being a real drift in circadian rhythm (**figure S6**). Thus, altogether, it appears that the deletion of *Ncaph2* and the absence of condensin II activity had an extremely limited effect on gene expression in adult hepatocytes.

**Table 1.**
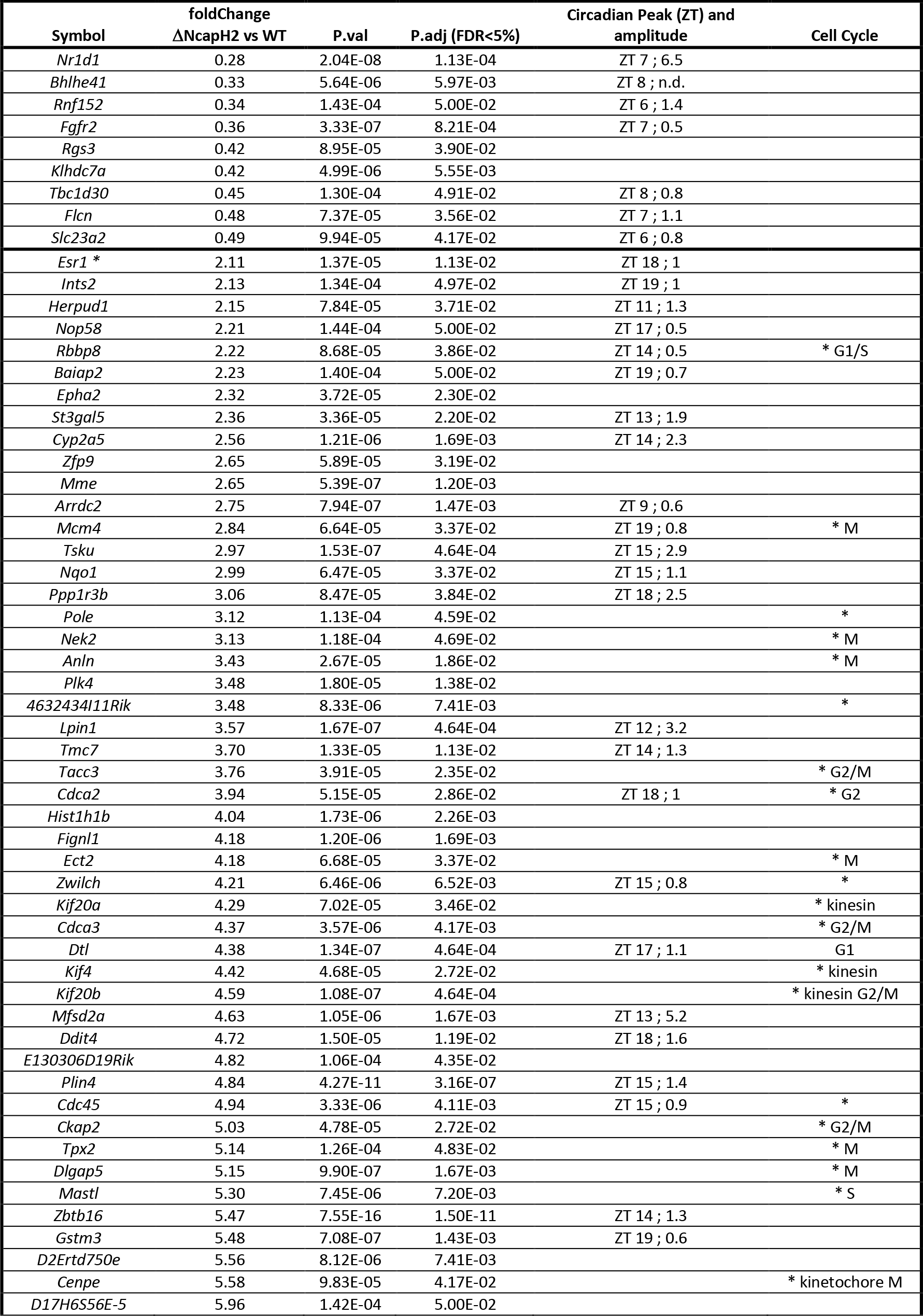
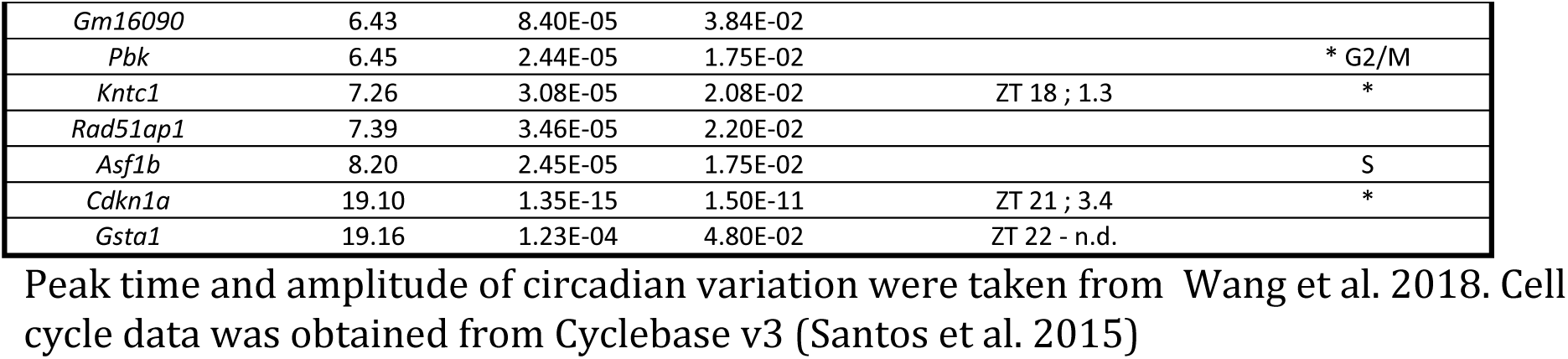
Differentially expressed genes in ΔNcaph2 samples Peak time and amplitude of circadian variation were taken from Wang et al. 2018. Cell cycle data was obtained from Cyclebase v3 (Santos et al. 2015)

**Table 2.** Molecular and cellular processes identified by GO-term enrichment analysis.

| GO term | Description | P-value | FDR q-value | Enrichment |
| --- | --- | --- | --- | --- |
| GO:0007049 | cell cycle | 1.38E-14 | 2.04E-10 | 7.79 (22) |
| GO:1903047 | mitotic cell cycle process | 3.07E-13 | 2.28E-09 | 10.29(17) |
| GO:0022402 | cell cycle process | 1.02E-12 | 5.03E-09 | 7.29 (20) |
| GO:0051301 | cell division | 2.98E-12 | 1.10E-08 | 9.92 (16) |
| GO:0051988 | regulation of attachment of spindle microtubules to kinetochore | 7.73E-08 | 1.43E-04 | 90.21 (4) |
| GO:0007051 | spindle organization | 7.77E-6 | 9.6E-3 | 12.61 (6) |

**Figure 4:**
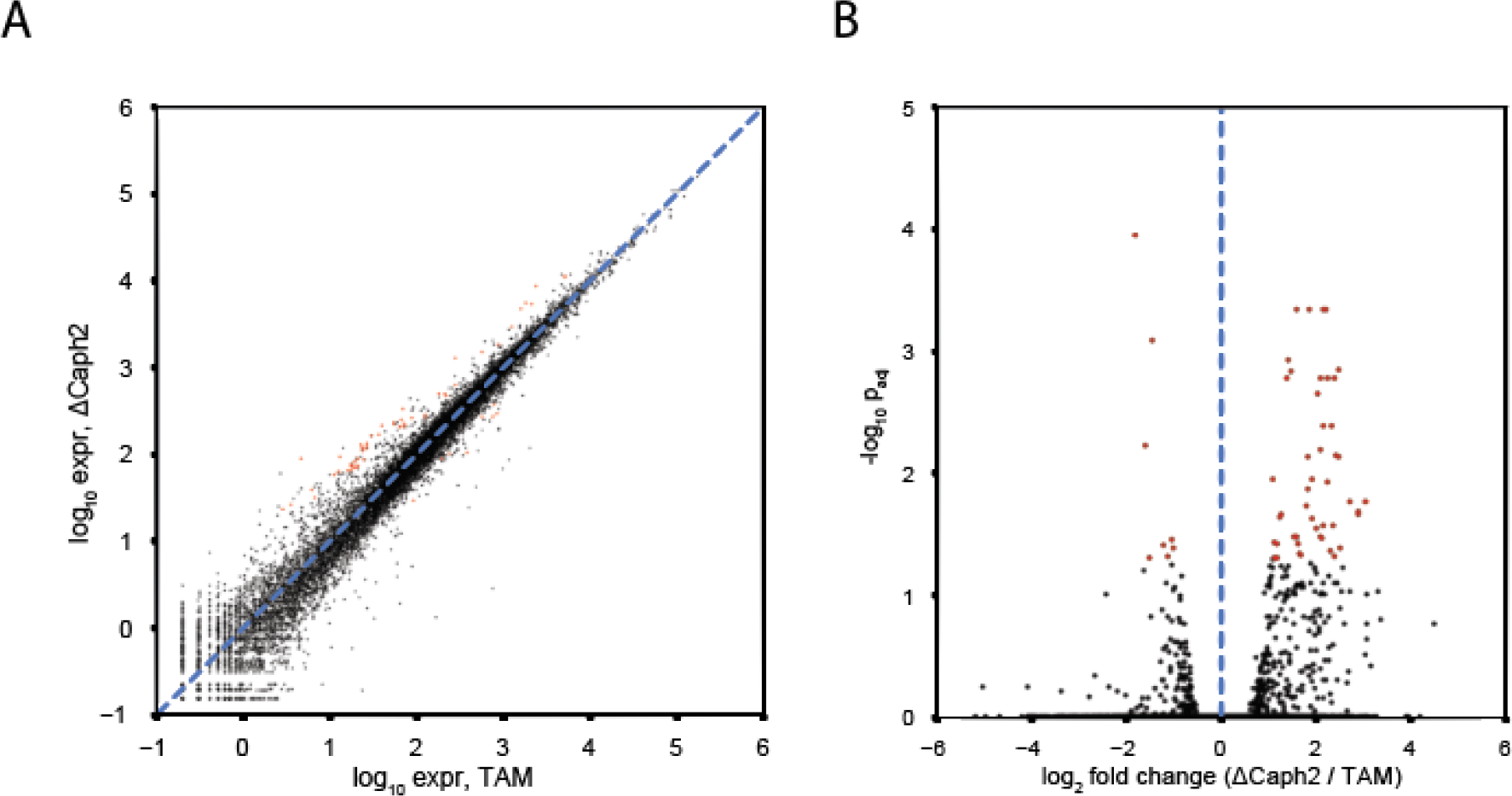
Condensin II removal has negligible effects on interphase gene expression. (A) Scatter plot of log expression (read counts normalized for sequence depth) between TAM and ΔNcaph2 conditions. (B) Volcano plot of differential expression analysis. Significantly differentially expressed genes at a false discovery rate of ≤ 0.05 are colored in red.

The absence of noticeable effects of a deletion of *Ncaph2* in non-dividing adult hepatocytes, as shown here, indicates that condensin II has no major function in maintaining interphase chromosome organization and gene regulation. One can argue that condensin I complexes - assuming that they are present in sufficient number in interphase nuclei - may compensate for the absence of condensin II in interphase. While we cannot formally exclude this hypothesis, our conclusion on the dispensability of condensin for interphase chromosomal organization is strengthened by a recent study in chicken DT40 cells, which reported major changes in mitotic chromosomes, but no significant differences the interphase chromosomes after acute degradation of either SMC2 or NCAPH2 in G2 phase (Gibcus et al., 2018) (re-analyzed in **figure 5**).

**Figure 5:**
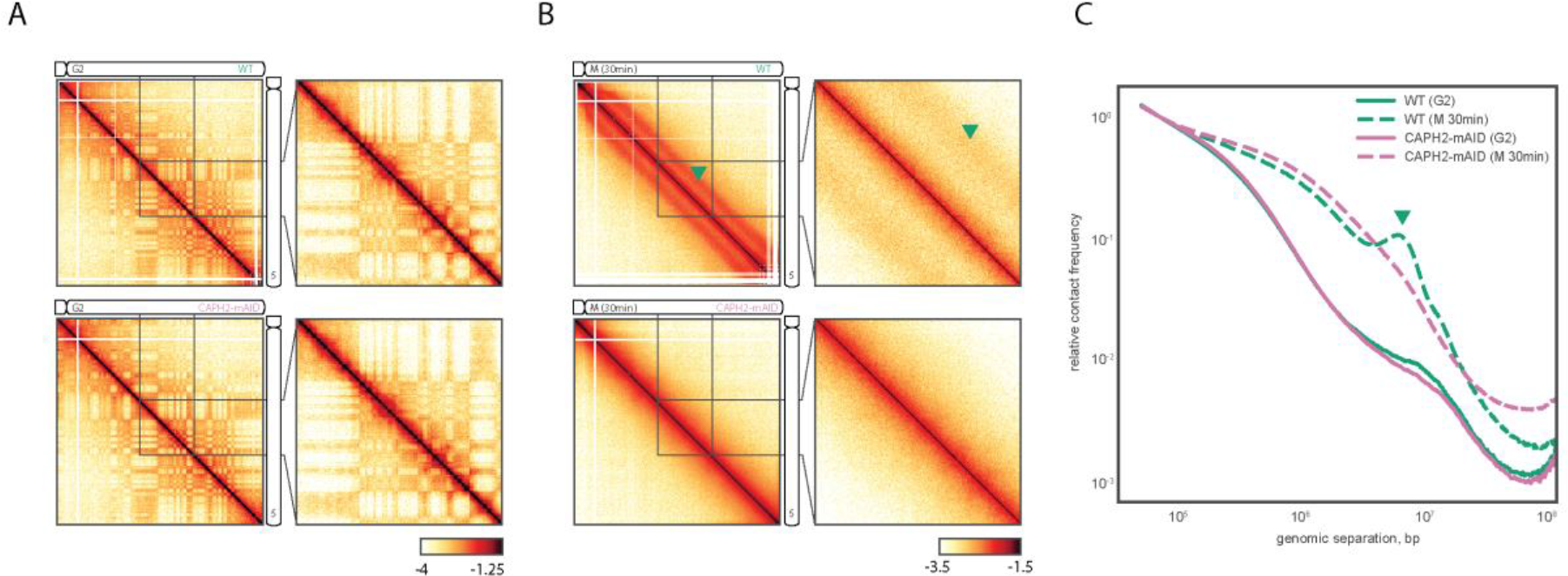
Condensin II degradation in synchronized DT40 chicken cells reveals severe impact on mitotic chromosome folding but no effect during G2. **(A)** Hi-C map of chromosome 5 in G2-arrested DK40 chicken cells in WT, above, and AID-mediated degradation of CAPH2, below. **(B)**Hi-C map of same regions in late prometaphase (t=30min). Top: WT, Bottom: CAPH2 depletion. **(C)**Genome-wide *P(s)* curves for the conditions in A and B, at both cell cycle stages. Data re-analyzed from Gibcus *et al*, 2018. Note the disappearance of the “bump” in the *P(s)* curve in the late prometaphase CAPH2 depletion condition (dashed curves). On HiC maps from cells in late prometaphase, this bump corresponds to a second diagonal band (green arrows in B), which disappears in the CAPH2 depletion condition. This is interaction pattern is indicative of a condensin II-dependent regular helical winding of a condensed chromatin scaffold (Gibcus et al, 2018). By contrast, CAPH2 depletion does not significantly affect *P(s)* or Hi-C maps in G2-arrested DK40 cells (solid curves in **C**, Hi-C maps in **A**).

Our findings that condensin II is largely dispensable for interphase gene expression and does not contribute noticeably to the maintenance of the different folding features characteristic of interphase chromatin contrast with several previous reports which argued that condensin does play an important role in the regulation of interphase mammalian chromosomal architecture (Dowen et al., 2013; Li et al., 2015; Yuen et al., 2017). However, the experiments in those studies were performed under very different conditions. Adult hepatocytes -- the system used here -- are slow-dividing and long-lived, producing perhaps two cell divisions over a period of 300 days (Magami et al., 2002). By contrast, the previous analyses of condensin function in mammalian cells were carried out using rapidly dividing cultured cell lines. Two of these studies ((Dowen et al., 2013; Yuen et al., 2017)) used mouse embryonic stem cells grown under standard culture conditions, which typically have a 10-12 hour-long cell division-cycle (Coronado et al., 2013), and the effects of knockdown experiments were assessed 3-5 days after knockdown. In the third study (Li et al., 2015), *NCAPH2* knock-down was initiated as MCF7 cells were progressively arrested from proliferation by serum deprivation, and its consequences were analyzed four days later. Even though this led to a final situation with 80-95% of the cell population in G0/G1 (Li et al., 2015), with a cell cycle length of approximately 20h (Cos et al., 1996) the majority of the originally asynchronous cell population would have completed one to three cell cycles before arresting at the mitogenic restriction point. Thus, these experimental designs leave open the possibility that the observed changes may result from defective mitotic condensation and segregation.

Condensin plays an essential role in mitotic chromosome assembly and segregation. Chromosomes are not correctly segregated if they exit mitosis as intermingled and potentially damaged structures, which can lead to global chromosomal instability and DNA damage responses (Ganem and Pellman, 2012), thus impacting gene expression and chromosome organization in the subsequent interphase. Consistent with this, recent studies in yeast also suggest that the regulatory changes associated with condensin deletion stem from indirect consequences of chromosomal instability rather than from a direct effect of condensin on transcription (Hocquet et al., 2018). Furthermore, in vertebrate cells, condensin complexes have been shown to occupy previously active promoters during mitosis, and this was proposed to “bookmark” them to allow efficient re-expression of the corresponding genes in G1 (Kim et al., 2013). Such ripple effects of mitotic perturbations may account for the previously observed consequences of condensin depletion in interphase cells, while they cannot impact our non- (or very rarely) dividing hepatocytes.

In addition to consequences of chromosome mis-segregation, the axial compaction of chromosomes initiated by condensin II and the lateral compaction subsequently reinforced by condensin I define a very specific organization, wherein each chromosome emerges at the end of mitosis as an individualized entity (Gibcus et al., 2018). Such an initial configuration may be part of the necessary steps for proper establishment of intra- and inter-chromosomal modes of organization in interphase. Accordingly, chromosomal territories (Bauer et al., 2012; Rosin et al., 2018), and at a smaller scale *cis*-interactions between highly active gene clusters (Yuen et al., 2017) and super-enhancers (Dowen et al., 2013), which have been shown to dependent on condensin activity, may be compromised if chromosomes exit mitosis as partially decondensed, intermingled structures as seen in condensin II-depletion mitosis (Gibcus et al., 2018; Hirano, 2016). Our present data argue that this dynamic organization, once established after cell division, does not require condensin II for its maintenance.

Altogether, our data provide evidence that contrarily to what has been previously suggested, condensin II complexes are dispensable to the maintenance of the different organizational modes adopted by interphase chromosomes in vertebrates. Instead, most of the effects of condensin depletion previously reported in “interphase” mammalian cells regarding chromosomal 3D interactions and gene regulation may stem from the indirect consequences of defective mitosis.

## Acknowledgments

We thank the EMBL Laboratory Animal Resources Facility for animal husbandry and the EMBL Genomics Core Facility for genomic library sequencing. We thank EUCOMM and the Wellcome Trust Sanger Institute for generously providing the Ncaph2 conditional mutant line. We thank John Marioni and Nuno Fonseca (EMBL/EBI) for advice with RNA-Seq data processing. W.S. and A.P were supported by an EMBL Interdisciplinary Postdoc (EIPOD) fellowship, under the Marie Sklodowska Curie Actions COFUND program. This work was partially supported by NIH (GM114190) and NSF,and by the Center for 3D Structure and Physics of the Genome of NIH 4DN Consortium (DK107980) to LM, and NIH Common Fund Program, grant U01CA200147, as a Transformative Collaborative Project Award to LM and FS. Work in the Spitz lab was supported by the European Molecular Biology Laboratory, the Institut Pasteur and the Deutsche Forschunggemeinschaft (DFG SP 1331/3-1). Funding from the European Commission’s seventh framework program through the collaborative research project RADIANT (Grant # 305626, to W.H.) contributed partially to this work.

## Author Contributions

W.S., C.H.H. and F.S. conceived the study and designed the experiments. W.S. generated experimental data with the help of A.P and I.A.S. W.H. provided resources and advice in preliminary data analysis. N.A. performed all computational analysis. L.M. provided advice and guidance on data analysis. N.A., W.S, C.H.H. and L.M. and F.S. analyzed the results and wrote the paper with input from the other authors.

## Material and methods

### Mouse lines

B6N;B6N-Ncaph2^<tm1a(EUCOMM)Wtsi>/Wtsi^ mice (thereafter referred as *Ncaph2* ^*lox/+*^) mice were generously provided by WTSI and maintained on C57BL/6J genetic background.

For deletion of the *floxed* exon and generation of ΔNcaph2 hepatocytes, *Ncaph2* ^*lox/+*^ were bred to *Ttr-cre/Esr1* (Tannour-Louet et al., 2002) transgenic mice. 12 week-old mice were injected with 1mg Tamoxifen (100μl of 10mg/ml Tamoxifen in corn oil) on 5 consecutive days. After another 5 days without injection, these mice were sacrificed and the hepatocytes were harvested as previously described (Schwarzer et al., 2017). Mice were genotyped by PCR using specific primer pairs (details available on request). Mouse experiments were conducted in accordance with the principles and guidelines in place at European Molecular Biology Laboratory, as defined and overseen by its Institutional Animal Care and Use Committee, in accordance with the European Directive 2010/63/EU.

### Western blots

Nuclear proteins from tamoxifen-treated wild-type C57BL/6J (TAM) and ΔNcaph2 hepatocytes were obtained by 10 min hypotonic lysis on ice in 10 mM HEPES-KOH pH 7.5, 10 mM KCl, 0.1 mM EDTA, 0.5% v/v NP-40, 1 mM DTT, 0.5 mM PMSF and Roche Complete protease inhibitor cocktail, followed by collection of the nuclei (5 min at 1000 *g*) and protein extraction in same buffer supplemented with 1% w/v SDS and 150 mM NaCl at 65 °C for 10 min. Total protein concentrations were estimated by Bradford assay (Biorad) and 11 μg protein was resolved by SDS-PAGE on 10% polyacrylamide or 3-8% NuPAGE Tris-Acetate (Thermo Fisher) gels and transferred to nitrocellulose. Membranes were pretreated and probed in PBS, 0.1% v/v Tween-20, 4% w/v non-fat milk powder with rabbit primary antibodies and goat anti-rabbit IgG-HRP (Dianova) secondary antibody.

As commercial antibodies raised against fragments of human NCAPH2 (Bethyl A302-275A; Abcam 83848) did not detect protein in the expected range, we raised a rabbit polyclonal antibody against full-length mouse NCAPH2. The antibody recognized multiple proteins from the mouse hepatocyte nuclear extracts, including a ~105 kDa protein that was depleted in ΔNcaph2 mice as well as human NCAPH2 in HeLa whole cell extracts.

### RT-qPCR

Unfixed hepatocyte aliquots were thawed and RNA was prepared with Qiagen RNeasy Kit. cDNA was generated using NEB ProtoScript^®^ First Strand cDNA Synthesis Kit with random primer mix. RT-qPCR was performed with Applied SYBR Green PCR Master Mix and following primers: Ncaph2-ex8/9-qPCR_F ACGGGAGTCCTGTTCCTGTA, Ncaph2-ex8/9-qPCR_R CTCTGCATCCTCCTCTCCAC, Gapdh-qPCR_F CTCCCACTCTTCCACCTTCG, Gapdh-qPCR_RCCACCACCCTGTTGCTGTAG, RTqPCR_Pgk1_Fwd TGGTATACCTGCTGGCTGGATGG and RTqPCR_Pgk1_Rev GACCCACAGCCTCGGCATATTTC.

### RNA-seq libraries and sequencing

Unfixed hepatocyte aliquots were thawed and RNA was prepared with Qiagen RNeasy Kit. RNA integrity was tested with Bioanalyzer (Agilent RNA Nano Kit) and ribosomal RNA was removed using Ribo-Zero rRNA Removal Kit (Illumina) prior to library preparation. Strand-specific libraries were prepared with NEBNext^®^ Ultra™ Directional RNA Library Prep Kit for Illumina^®^. After amplification and size selection with Agencourt AMPure XP beads (Beckmann Coulter) their size-distributions were determined with Bioanalyzer. Equimolar pools of libraries were sequenced with Illumina HiSeq2000 (50bp, single end). Reads were uniquely mapped to the reference genome NCBI37/mm9.

### Tethered Chromatin Capture (TCC)

Roughly 100 mio fixed hepatocytes per sample were processed according to Kalhor et al. (Kalhor et al., 2012) using HindIII. Libraries were PCR-amplified (12 cycles) and size selected with E-Gel^®^ SizeSelect™ (Thermo Fisher). Equimolar pools of libraries were sequenced with Illumina HiSeq2000 (50bp, paired end). We retrieved between 100 and 150 mio paired reads per sample, of which ~40% had both sides uniquely mapped to the reference genome NCBI37/mm9.

### Computational Analysis

#### Hi-C data processing

Hi-C datasets were processed using a portable pipeline driven by the *nextflow* workflow manager (Di Tommaso et al., 2017) called *distiller*, available at https://github.com/mirnylab/distiller-nf. Briefly, we mapped Hi-C sequencing reads to the mouse reference assembly mm9 using *bwa mem* (Li 2013) with flags -SP. Alignments were parsed, filtered for duplicates and pairs were classified using the *pairtools* package (https://github.com/mirnylab/pairtools). Hi-C pairs were aggregated into contact matrices in the cooler format using the *cooler* package (https://github.com/mirnylab/cooler) at multiple resolutions. All contact matrices were normalized using the iterative correction procedure (Imakaev et al., 2012) after bin-level filtering. Low-coverage bins were excluded using the MAD-max (maximum allowed median absolute deviation) filter on genomic coverage, described in (Schwarzer et al., 2017), using a threshold of 5.0 MADs. To remove short-range Hi-C artifacts-such as unligated and self-ligated Hi-C molecules-we removed contacts mapping to the same or adjacent genomic bins, up to a distance of at least 10kb. The same procedure was used to process the DT40 Hi-C data sets from (Gibcus et al., 2018). Expected contact profiles were obtained by dividing each diagonal of a *cis* contact matrice by its average value over non-filtered genomic bins. This was performed using the *cooltools* package (https://github.com/mirnylab/cooltools).

#### P(s) curves

Scaling curves of contact frequency P as a function of genomic separation *s* were generated using *cooltools* by aggregating normalized contact frequency over valid pixels along diagonals of 1kb-resolution *cis* contact maps, with diagonals grouped into geometrically increasing spans of genomic separation. Average contact frequency P(s) curves are displayed using log-log axes.

#### Compartmentalization analysis

Compartmentalization as reflected in Hi-C maps was quantified using an eigenvector decomposition procedure based on (Imakaev et al., 2012), as implemented in the *cooltools* package. In short, compartment tracks (that is, the values or scores of compartment signal across all genomic bins) were quantified as the dominant eigenvector of the observed or expected 20-kb and 100-kb *cis* contacts maps upon subtraction of 1.0 and rescaling by the magnitude of the square root of its eigenvalue.

Compartmentalization strength was assessed by taking the average contact enrichment between pairs of loci with whose eigenvector scores lied in the extremes of the distribution of eigenvector scores and computing the ratio (AA + BB)/(2*AB), where AA refers to the average enrichment of loci whose two bins both lie in the top n% of eigenvector scores, BB to that of loci whose two bins both lie in the bottom n% of eigenvector scores, and AB to that of loci whose two bins have strong A and B compartment signal, respectively. The fraction at n=20% was used as an overall measure of compartmentalization strength. Saddle plot heatmaps and compartmentalization strength profiles were created using the *cooltools* package.

#### Domain detection and peak coordinates

We used the domains detected in the WT condition using in the *lavaburst* package (https://github.com/nvictus/lavaburst) from (Schwarzer et al., 2017). While domain callers do not distinguish compartmental domains from TADs, the majority of domains identified in the WT condition were shown to correspond to TADs. To characterize the structure of known Hi-C peaks in our data, we used the list of peaks detected in Hi-C maps for the CH12-LX mouse cell line (Rao et al., 2014).

#### Insulation analysis

Diamond insulation scores (Crane et al., 2015) were calculated on all 20-kb and 40-kb maps using the *cooltools* package. Additionally, an insulation minima calling procedure based on peak prominence, described in (Nora et al., 2017) was used to define insulating loci. We considered only insulating loci called in both replicate Hi-C maps and in the pooled Hi-C maps for each condition, at 20-kb resolution. Differential insulation analysis was performed using quantile-normalized log_2_ insulation scores on ~11,000 loci classified as insulating in at least one of the four conditions. Since each pair of treatment-control conditions considered (WT vs TAM, WT vs ΔNcaph2, WT vs ΔNipbl) consisted of two replicates each, we compared the log_2_ ratios of insulation score between conditions (treatment vs control) and repeated the same procedure on ratios within the same condition (replicate-1 vs replicate-2). Because our sample size for hypothesis testing on each locus was small (*n=2*), we use the regularized t-test (Baldi and Long, 2001; Cui and Churchill, 2003) that combines information from the locus-specific variance in log-ratio in with a background variance estimate obtained from the bulk of insulating sites considered. We use a positive false discovery rate (Storey, 2003) of 0.05 on all three sets of hypothesis combined to set a common significance threshold.

#### ChIP-seq processing and H3K27ac clustering

We used H3K27ac ChIP-seq peaks and motif assignments from (Schwarzer et al., 2017). To identify regions that exhibit enhancer characteristics, we followed the H3K27ac peak-stitching strategy using a 12.5kb stitching distance as in (Hnisz et al., 2013). Rather than defining a cutoff for super-enhancer status, we divided the stitched regions into equal sized groups, first by length, then by expression level as quantified by RNA-seq coverage in WT from (Schwarzer et al., 2017). Additionally, we obtained a curated list of super-enhancers for mouse liver from dbSuper (Khan and Zhang, 2016). ChIP-seq data from (Canella et al., 2012)and (Yuen et al., 2017) were reprocessed following the steps of the ENCODE ChIP-seq pipeline (https://github.com/ENCODE-DCC/chip-seq-pipeline). Peaks were also divided into two equal sized groups by expression level.

#### Hi-C pileup maps

We used *cooltools* to calculate aggregate iteratively corrected or observed-over-expected contact frequency maps (pileup maps) centered at specific genomic loci and bounded by a fixed flanking genomic distance. For single genomic landmarks or intervals (TAD/peak calls, CTCF sites), pileup maps are centered on the main diagonal at each feature’s midpoint. For the Hi-C pileup maps of paired genomic loci centered away from the Hi-C map’s main diagonal, we selected 5000 random pairs without replacement in *trans* and for each range of genomic distances in *cis* from each set of landmark features. We then used *cooltools* to calculate observed-over-expected Hi-C pileup maps centered at the midpoint coordinates of the landmark feature pairs.

#### RNA-seq and GO enrichment analysis

We mapped the RNA-seq data to the mm9 reference mouse genome assembly and GENCODE vM1 transcriptome using *STAR* v.2.5.0a60 (Dobin et al., 2013). To obtain the tracks of local transcription, we aggregated the uniquely mapped reads into RPM-normalized bigWig files using the built-in *STAR* functionality. To find differentially expressed genes, we aggregated the read counts at the gene level using HTSeq (Anders et al., 2015) with the ‘union’ option and called differentially expressed genes with DESeq2 (Love et al., 2014).

#### Circadian rhythm and cell cycle data analysis

We used the data from (Wang et al., 2018), except for *Gsta1* (Zhang et al., 2009) and *Bhlhe41* (Tognini et al., 2017). Cell cycle information was inferred from (Santos et al., 2015).

## Supplementary Figures

**Figure S1.**
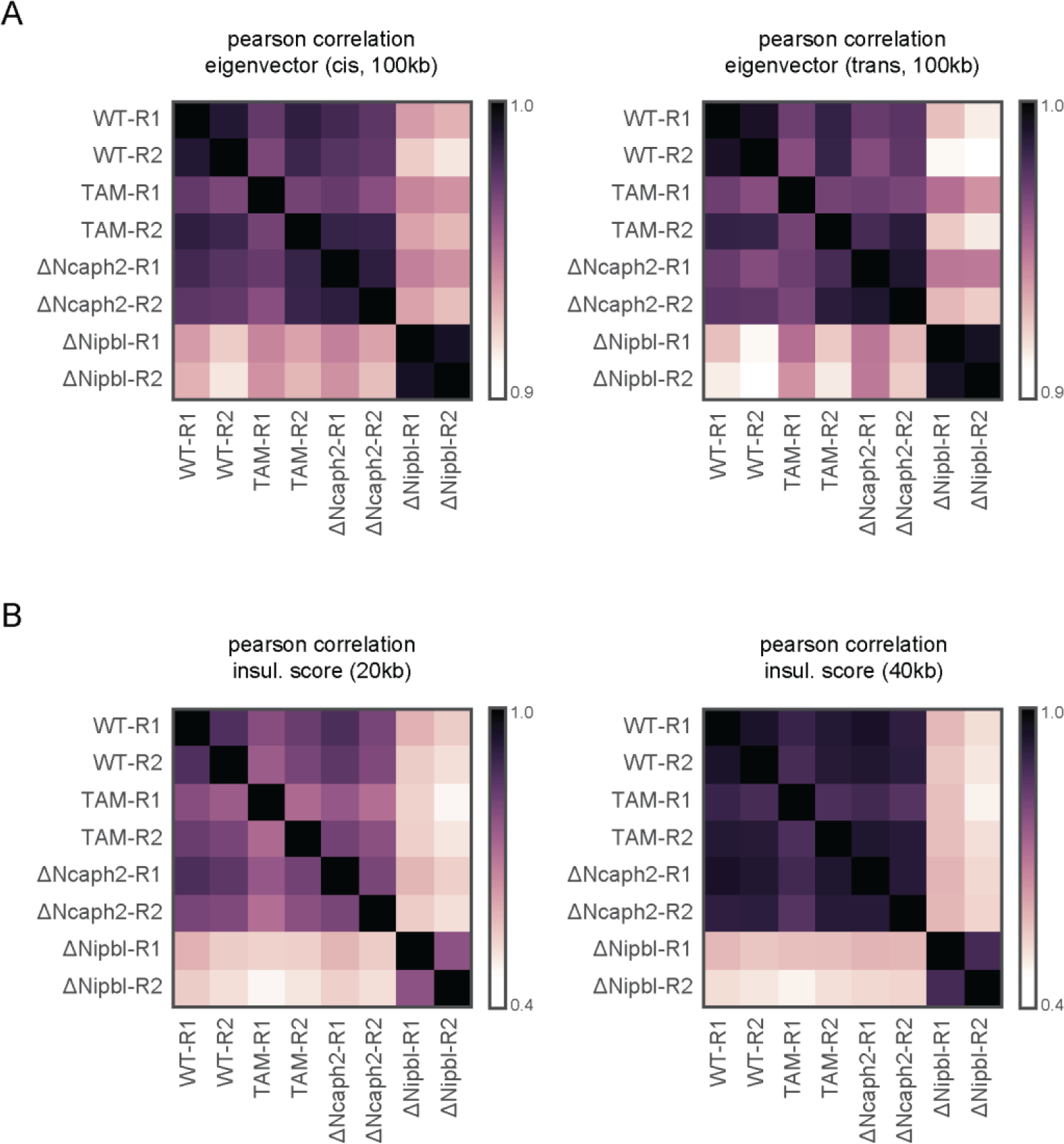
Replicates. Pearson correlation scores comparing the different TTC experimental conditions and replicates for insulation score (A) eigenvector signal extracted from cis (left) and trans (right) at 100kb resolution and (B) insulation score at 20kb (left) and 40kb (right) resolution. WT = untreated control (*Nipbl* ^*flox/flox*^; *Tg(Ttr-cre/Esr1*)*^*1Vco*^; no induction of Cre by tamoxifen injection). TAM = tamoxifen-injected control animals (*Ncaph2*^*flox/flox*^; TAM-injected; animals without *Tg(Ttr-cre/Esr1*)*). ΔNcaph2 = Conditional Ncaph2 deletion (*Ncaph2*^*flox/flox*^; *Tg(Ttr-cre/Esr1*)*^*1Vco*^−TAM-injected). ΔNipbl = Conditional Nipbl deletion (*Nipbl*^*flox/flox*^; *Tg(Ttr-cre/Esr1*)*^*1Vco*^−TAM-injected) (Schwarzer et al., 2017).

**Figure S2.**
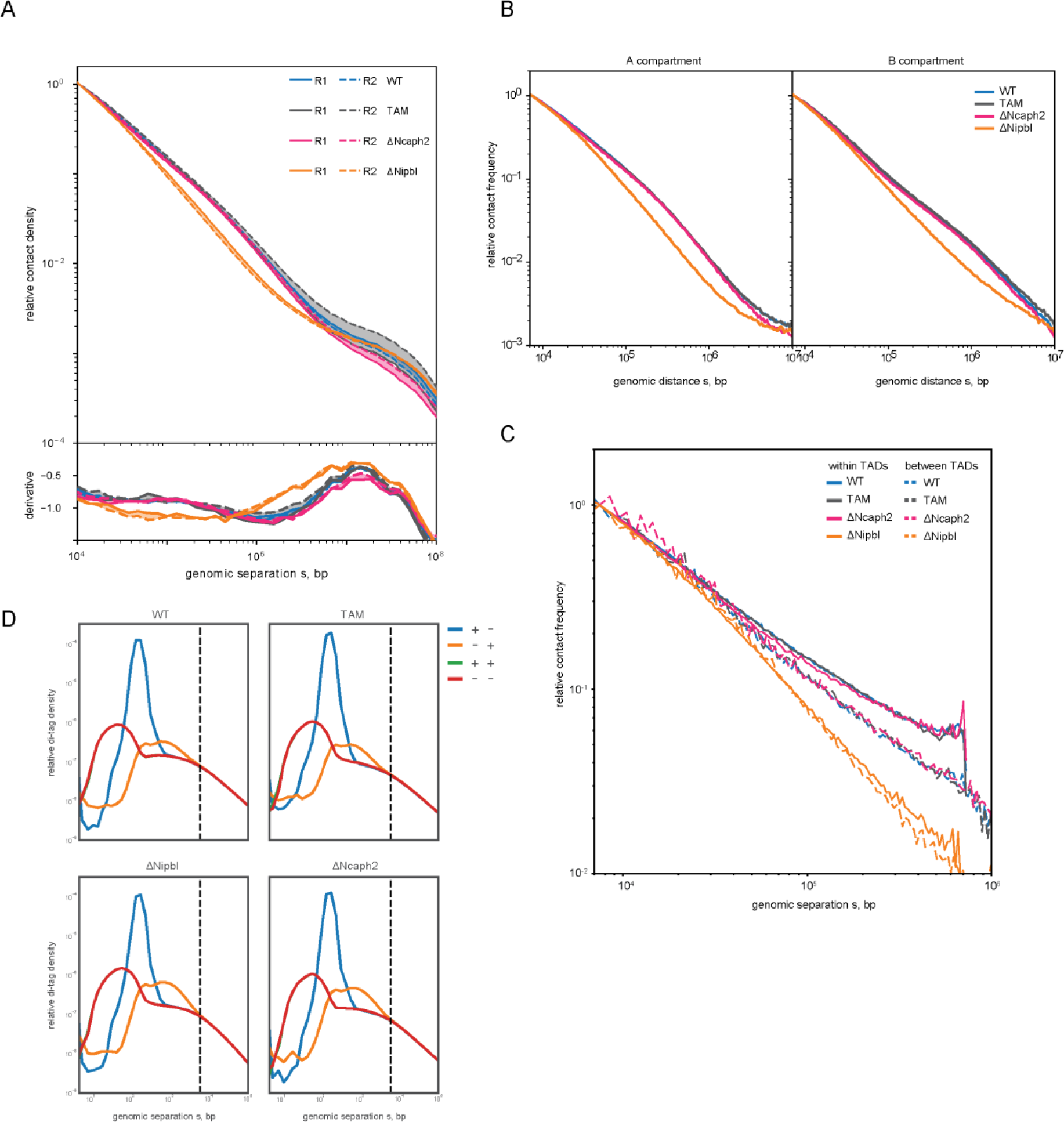
P(s) curves. *P(s)* curves by replicate and in sub-regions. (A) *P(s)* curves as in Figure 2a but calculated separately for each replicate. The areas between curves from the same condition are shaded. (B) *P(s)* curves for each condition, restricted to regions within A compartmental intervals (left) and B compartmental intervals (right). The curves for ΔNCaph2 and controls follow the same trend, and the shoulder in B intervals is slightly more right-shifted (implying larger loop length in B regions). However, in both cases the ΔNipbl shoulder is lost. (C) *P(s)* curves within and between TADs called in WT. As observed in (Schwarzer et al., 2017), the shallower decay of the intra-TAD contact frequency scaling is lost in ΔNipbl and the two curves collapse. On the other hand, ΔNCaph2 curves match those of WT and TAM. (D) Short distance scaling curves of mapped read pairs split by strand orientations. The divergence of the curves reflects contaminant ligation products of the Hi-C protocol. The point of convergence is used to determine a threshold for short distance filtering.

**Figure S3.**
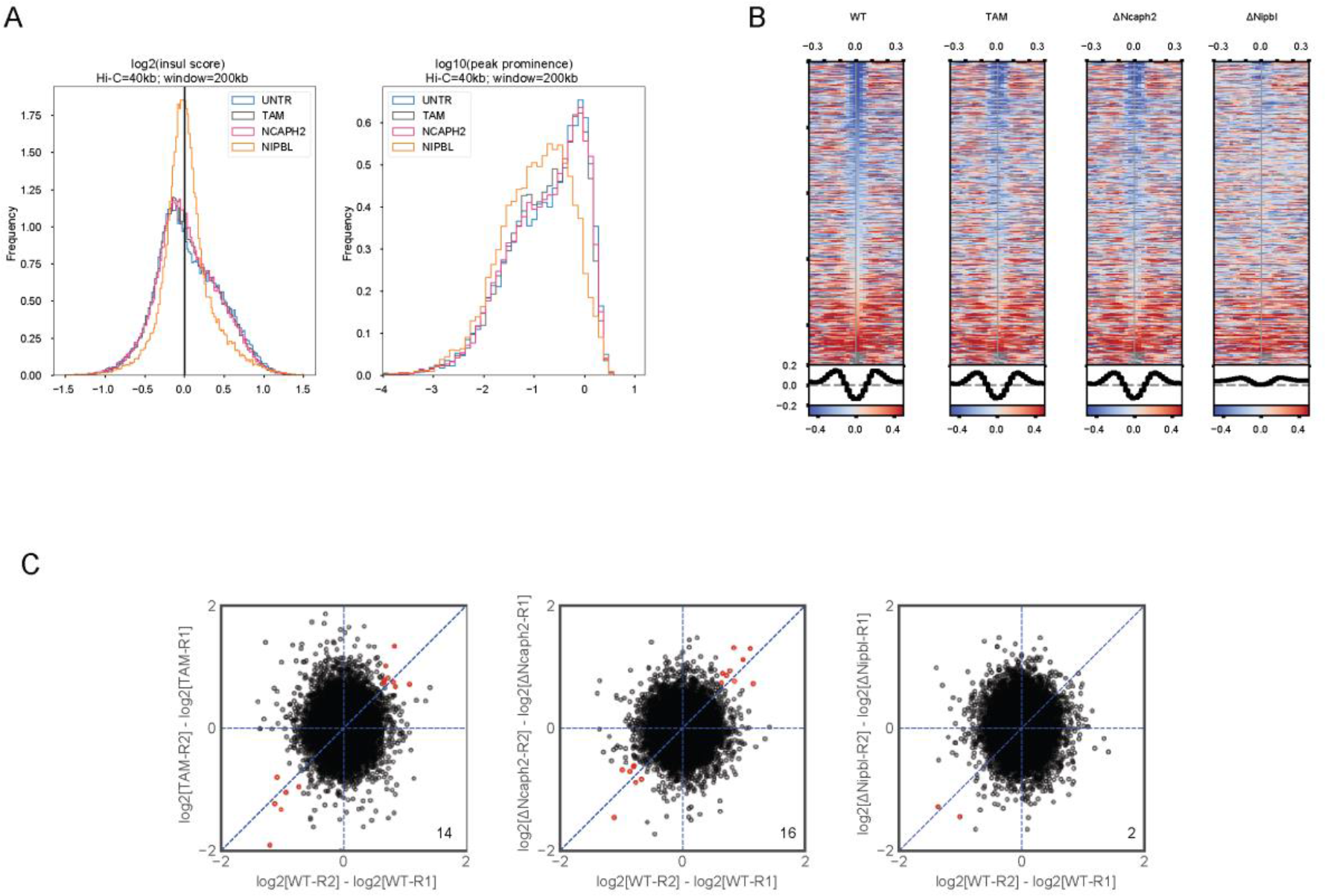
Insulation analysis. (A) Histograms of log_2_ insulation score (left) and log_10_ peak prominence (right). (B) Stacked heatmap of insulation scores at wildtype insulation valleys (as in figure2k). (C) As a control for differential insulation analysis, we compare intra-condition log fold changes of replicates using the same background variance estimate. The number of hits is printed in the bottom right corner.

**Figure S4.**
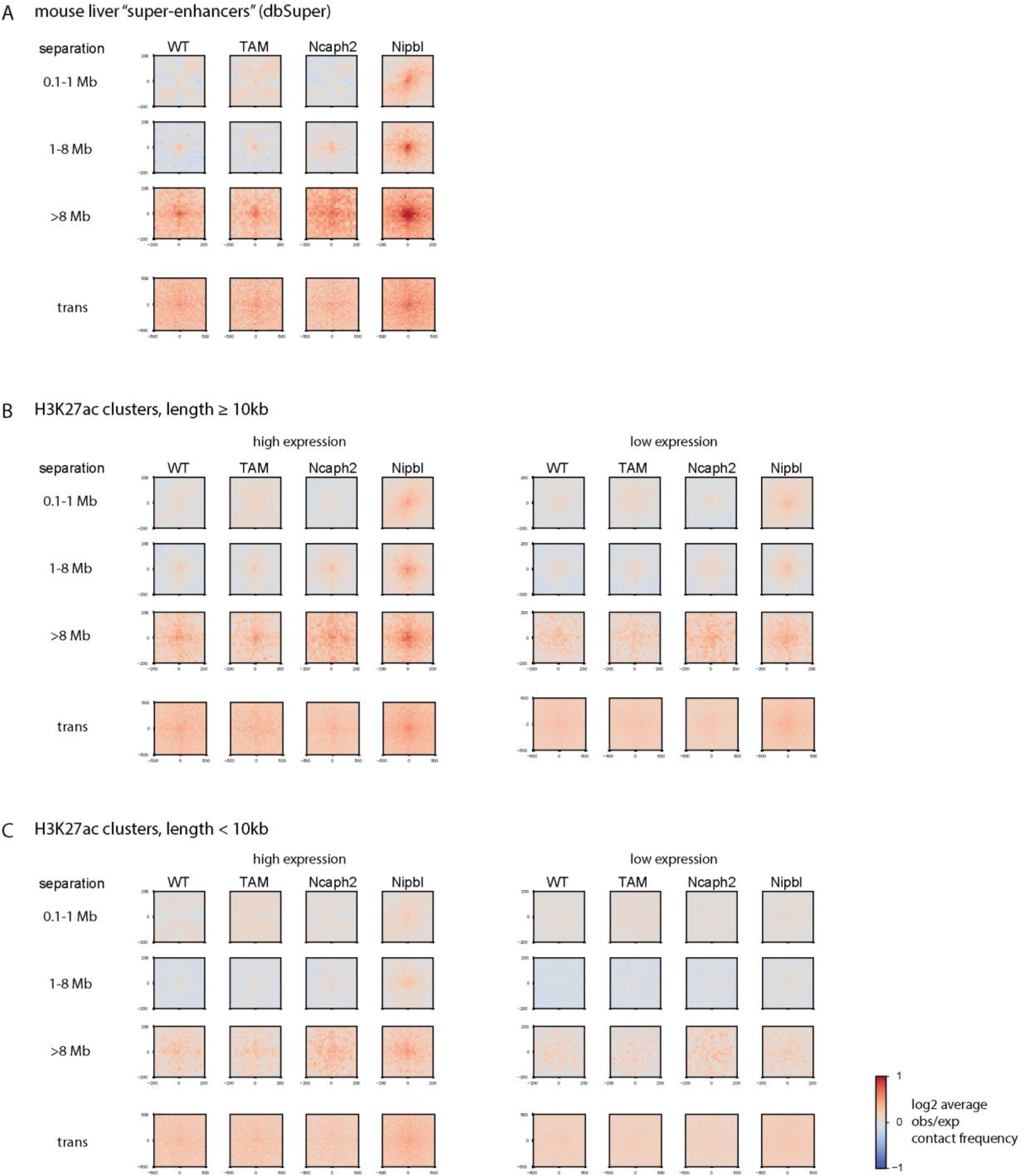
Aggregate paired landmarks (super-enhancers) Aggregate Hi-C pileup maps of pairs of super-enhancers stratified by genomic separation in cis and in trans. (B) Aggregate Hi-C pileup maps of pairs of long, stitched H3K27ac peaks not selected for super-enhancer status, but rather divided into two equal sized expression groups by the mean RNA-seq coverage measured in WT. (C) Same, but for the remaining unstitched H3K27ac peaks.

**Figure S5.**
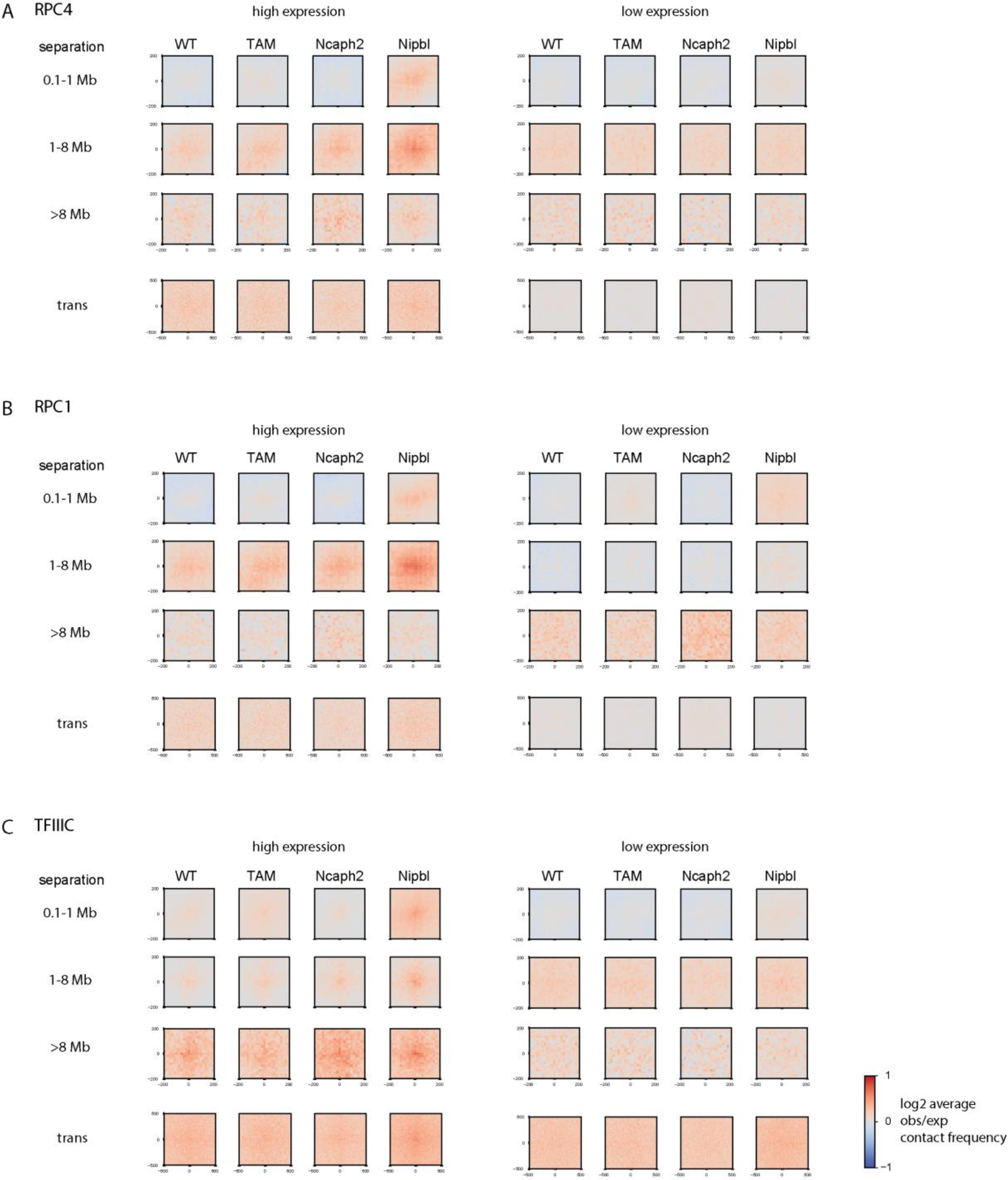
Aggregate paired landmarks (Pol3, TFIIIC) Aggregate Hi-C pileup maps of genomic landmarks stratified by genomic separation in *cis* and in *trans*: pairs of binding sites for RNAP-III associated factors (A) RPC4/POLR3D and (B) RPC1/POLR3A from mouse liver (Canella et al., 2012); (C) pairs of binding sites for TFIIIC measured in mouse ES cells (Yuen et al., 2017). Loci were divided into two equal sized expression groups by the RNA-seq coverage measured in WT.

**Figure S6.**
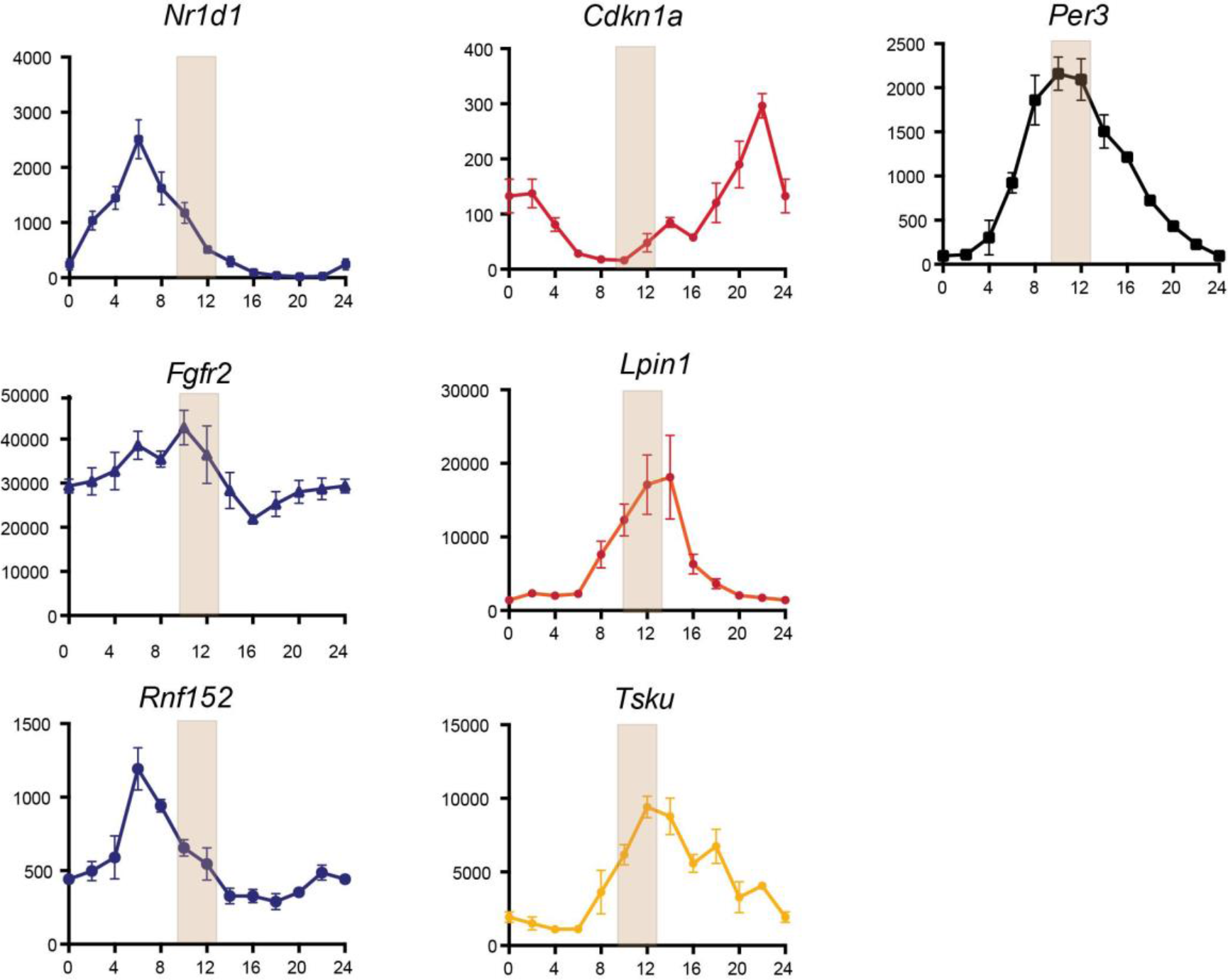
Circadian expression of several genes with expression differences between ΔNcaph2 and TAM controls. Relative expression levels are plotted over a 24h-period. In blue (resp. red/orange), genes which are downregulated (resp. up-regulated) in ΔNcaph2 samples. Down-regulated genes correspond to early-peak (ZT=6-8), while up-regulated ones peak latter in the day (ZT>15) (**see Table 1**). The brown shaded bar show the period during which samples have been collected. Circadian genes (e.g. Per3) which expression is stable during that period do not show expression changes between ΔNcaph2 and controls. Data from (Fang et al., 2014; Wang et al., 2018).

**Table S1.** Hi-C mapping statistics

| **sample** | **codename** | **raw read pairs** | **valid pairs** | **cis** | **trans** | **cis:total** |
| --- | --- | --- | --- | --- | --- | --- |
| WT-R1 | 014637Untr | 139,343,707 | 53,326,649 | 41,809,285 | 11,517,364 | 0.78 |
| WT-R2 | 019440Untr | 114,258,179 | 42,577,162 | 33,363,853 | 9,213,309 | 0.78 |
| WT |  |  | 98,775,892 | 75,173,138 | 23,602,754 | 0.76 |
| TAM-R1 | 014199TAMcontr | 94,032,309 | 14,065,226 | 12,075,375 | 1,989,851 | 0.86 |
| TAM-R2 | 014399TAMcontr | 134,045,253 | 51,685,051 | 39,445,882 | 12,239,169 | 0.76 |
| TAM |  |  | 67,529,373 | 51,521,257 | 16,008,116 | 0.76 |
| ΔNipbl-R1 | 014200Nipbl | 131,409,162 | 43,074,909 | 32,545,604 | 10,529,305 | 0.76 |
| ΔNipbl-R2 | 016350Nipbl | 133,914,046 | 50,686,600 | 38,098,292 | 12,588,308 | 0.75 |
| ΔNipbl | | | 95,471,646 | 70,643,896 | 24,827,750 | 0.74 |
| ΔNcaph2-R1 | 014398Ncaph2 | 127,595,875 | 42,011,634 | 31,569,049 | 10,442,585 | 0.75 |
| ΔNcaph2-R2 | 014953Ncaph2 | 116,070,234 | 42,168,415 | 29,788,754 | 12,379,661 | 0.71 |
| ΔNcaph2 | | | 86,316,825 | 61,357,803 | 24,959,022 | 0.71 |

